# The ability of *Pseudomonas aeruginosa* to adopt a <u>S</u>mall <u>C</u>olony <u>V</u>ariant (SCV) phenotype is conserved, and not restricted to clinical isolates

**DOI:** 10.1101/2021.02.05.430018

**Authors:** Alison Besse, Mylène Trottier, Marie-Christine Groleau, Eric Déziel

**Author notes:** Address correspondence to Eric Déziel.

## Abstract

A subpopulation of Small Colony Variants (SCVs) is a frequently observed feature of *Pseudomonas aeruginosa* isolated from cystic fibrosis (CF) lungs biofilms. SCVs have almost exclusively been reported from infected hosts, essentially CF individuals or, by extension, from laboratory cultivation of strains originated from infected hosts. We previously reported the identification of *P. aeruginosa* SCVs emerging from a non-clinical strain and displaying features shared with clinical SCVs. In the present work, we investigated the ability of 22 *P. aeruginosa* isolates from various environmental origins to, under laboratory culture conditions, spontaneously adopt a SCV-like smaller alternative morphotype distinguishable from the ancestral parent strain. Unexpectedly, we found that all the *P. aeruginosa* strains tested have the ability to adopt a SCV morphotype, regardless of their origin. Based on the phenotypes already described for SCVs, the SCV-like morphotypes obtained were clustered in two groups displaying various phenotypic profiles, including one characteristic of already described SCVs. We conclude that the ability to switch to a SCV phenotype is a conserved feature in *Pseudomonas aeruginosa*.

**IMPORTANCE:** *P. aeruginosa* is an opportunistic pathogen that thrives in many environments. It is significant public health concern, notably because it is the most prevalent pathogen found in the lungs of people with cystic fibrosis (CF). In infected hosts, its persistence is believed to be related to the emergence of an alternative small colony variant (SCV) phenotype. By reporting the distribution of *P. aeruginosa* SCVs in various non-clinical environments, this work contributes to understanding a conserved adaptation mechanism used by *P. aeruginosa* to rapidly adapt in all environments. Counteraction of this strategy could prevent *P. aeruginosa* persistent infection in the future.

## INTRODUCTION

The high genomic and metabolic diversity of *Pseudomonas aeruginosa* allows this bacterium to thrive in diverse environments, such as aquatic habitats, soil, food, and even built environments, such as hospital premise plumbing systems (1-3). This opportunistic pathogen, frequently identified as a causative agent of nosocomial infections, is a major cause of infections in immunocompromised individuals. Notably, *P. aeruginosa* is the most prevalent pathogen found in the lungs of people with cystic fibrosis (CF) (4-6).

*P. aeruginosa* expresses a broad range of virulence determinants that counteract the host immunity and promote survival (7). One of these factors is the ability to form biofilms. These organized communities largely contribute to evade host immunity and antimicrobial treatments. For instance, the biofilm matrix delays penetration of antibiotics and host defense effectors (8-10). *P. aeruginosa* typically persists in the lungs of CF individuals as a biofilm (11, 12).

The emergence of a subpopulation of Small Colony Variants (SCVs) is a frequently observed feature of *P. aeruginosa* isolates from CF lungs biofilms (13, 14). SCVs are characterized by circular opaque dwarf colonies with a diameter about three-time smaller than wild-type colonies (WT) (14-17). Shortly after their first report, we proposed that SCVs are phenotypic variants (18). Phenotypic variants arise from a phase variation mechanism, traditionally defined as a high-frequency ON/OFF switch between phenotypes in a heritable and reversible manner (19-21). Indeed, spontaneous reversion to the wildtype-like morphotype has been observed for SCVs (18, 22)

SCVs exhibit cell surface hyperpiliation and adherence to abiotic surfaces (16, 18, 23). These properties promote biofilm formation (24). Consistent with enhanced biofilm formation, a motility deficiency, notably flagellar, has also been observed for SCVs (16, 18, 25). Additionally, SCVs exhibit autoaggregative properties (16, 23). Many of these phenotypes are linked to an overproduction of exopolysaccharides (EPS) (alginate, Pel and Psl) by SCVs (14, 26). These phenotypes have in common to be regulated by the intracellular second messenger c-di-GMP though binding to specific receptors. For instance, high c-di-GMP levels activate the expression of the *pel* operon, leading to production of the EPS Pel, and repress flagellar motility (27-29).

It is striking that SCVs have almost exclusively been isolated from infected hosts, essentially CF individuals; or by extension, from laboratory cultivation of strains sampled from infected hosts (13). For instance, several studies have recovered SCVs from lung, sputum or deep throat swabs of CF individuals (12, 16, 17, 30). CF is not the only pathology associated with the emergence of *P. aeruginosa* SCVs. These variants have also been isolated from urine, feces, endotracheal secretion and pleural effusion of patients suffering from meningioma, anoxic encephalopathy, hepatocellular carcinoma, lung carcinoma or grave asphyxia neonatorum (31). While SCVs have been generated under *in vitro* and *in vivo* laboratory conditions, there emergence seems always associated with a clinical infection environment. For instance, SCVs have been generated *in vitro* in tube biofilms from the prototypic clinical strain *P. aeruginosa* PAO1 (23). SCVs have also been obtained *in vivo* from *P. aeruginosa* strains during infections in burn wound porcine models and murine models (32, 33).

Intriguingly, 20 years ago we reported one of the first identification of *P. aeruginosa* SCVs, that quickly emerged when a soil isolate was grown on a non-aqueous phase liquid, hexadecane, as sole substrate (18). The SCV morphotype of strain 57RP predominates when biofilm growth conditions are preferable and displays features shared with clinical SCVs: high adherence, efficient biofilm formation, hyperpiliation and reduced motility (18).

Since most SCVs have until now been isolated from clinical samples, it remains unclear how widespread is the ability of *P. aeruginosa* to exploit phase variation and develop this phenotype. In this work, we investigated the ability of *P. aeruginosa* isolates from various environmental origins to spontaneously adopt, under laboratory culture conditions, a SCV-like smaller colony morphotype readily distinguishable from their ancestral parent. We tested 22 *P. aeruginosa* strains from four different categories of environments: soil, food, hospital water systems and clinical. We found that all the *P. aeruginosa* strains have the ability to adopt the SCV phenotype, regardless of their origin.

## RESULTS

### The ability to form SCV-like morphotype colonies is a conserved feature in *Pseudomonas aeruginosa*

A few SCVs of *P. aeruginosa* have been reported under various culture conditions promoting biofilm formation (16, 18, 23). In order to more broadly investigate the ability of *P. aeruginosa* to adopt a SCV-like morphotype, we cultured 22 isolates from various origins in static liquid medium for 65 h then spread onto TSA plates to obtain isolated colonies. Six strains were from food samples (meat and fish from markets), six from clinical samples (5 from CF patients and the clinical prototypic strain PA14 from a burn patient), five from petroleum oil-contaminated soil and five from hospital sinks (drain, splash area and tap) (Table 1, columns 1 and 2). To cover the variety of temperatures relevant to these various habitats, the cultures were incubated in a temperature range varying from 30 to 40°C. At the onset, none of the strains were displaying a SCV phenotype (data not shown), but after 65 h of incubation all isolates diversified in a range of colony morphotypes, including small colonies that appear typical of SCVs (Fig. 1, for selected strain from each origins). Small colonies emerged in the cultures incubated at all tested temperatures (data not shown).

**Table 1.**
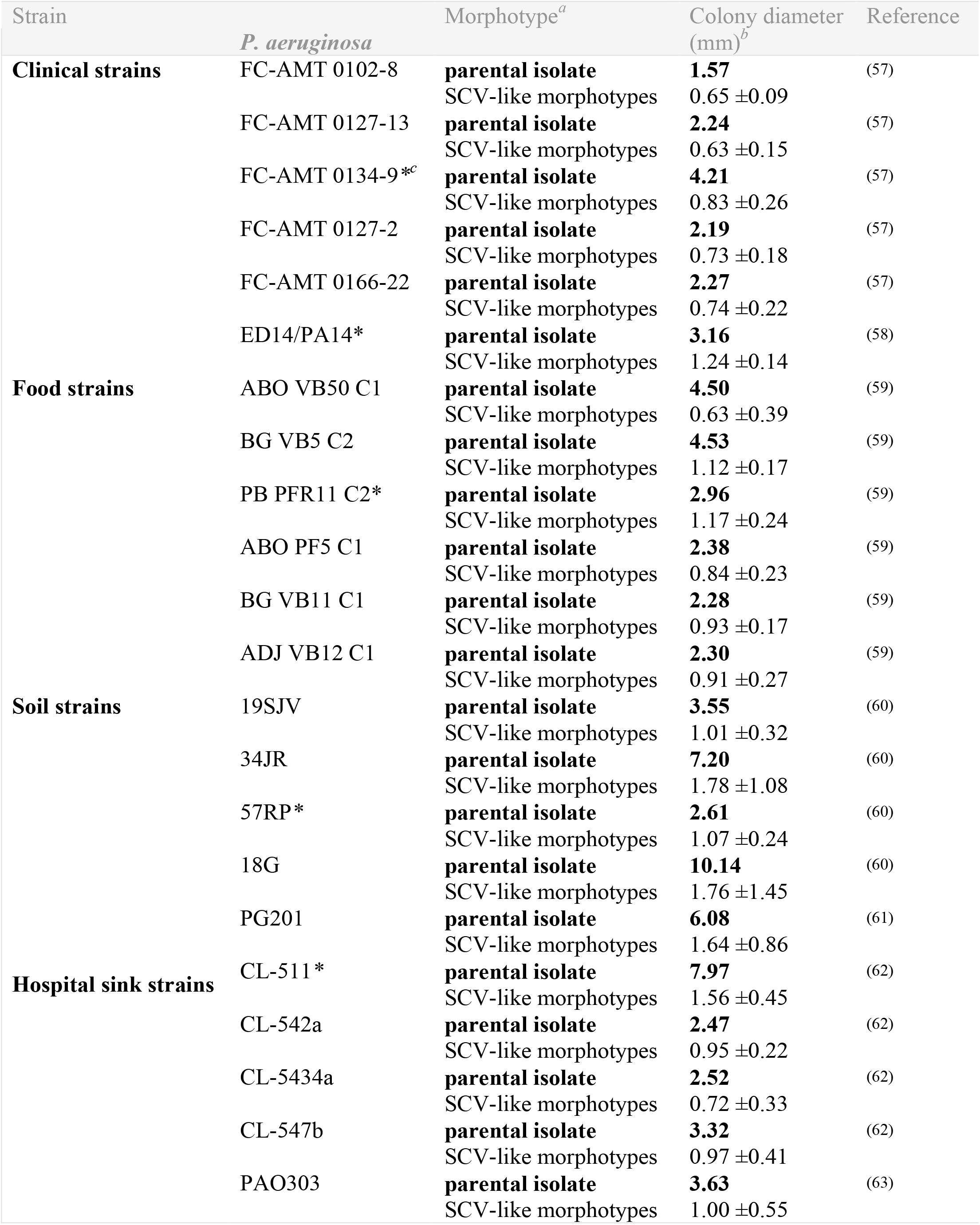
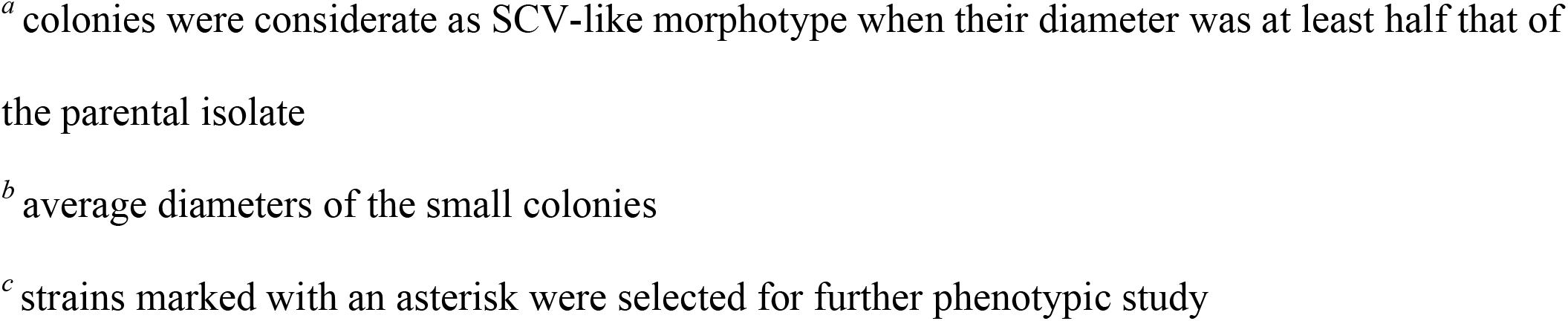
Colony diameters and phenotypes of parental isolates and their static liquid culture evolved small morphotypes.

**Fig 1.**
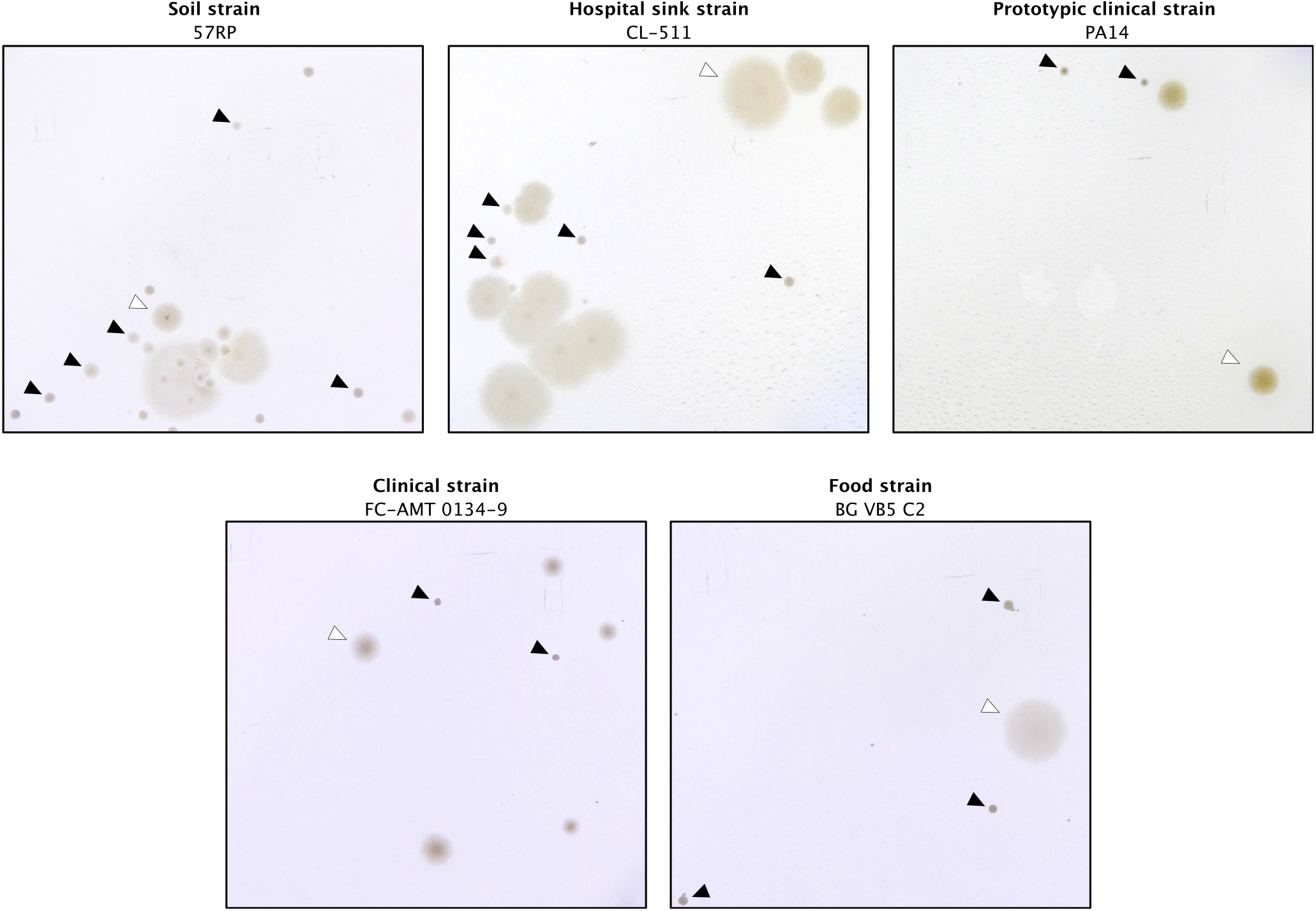
Small colonies of *Pseudomonas aeruginosa* emerge in static cultures from strains isolated from various origins. Parental strains were inoculated under static liquid conditions in TSB for 65 hours and spread onto TS-Agar 2% plates. Black arrows indicate smaller colonies. White arrows indicate parent-like colony.

Reported SCVs have an average diameter two to four times smaller than WT colonies. Colonies correspondingly smaller than the parental strains emerged from all 22 strains (Table 1). This result strongly suggests that the ability to produce variant colonies displaying an SCV-like morphotype is a conserved feature of *P. aeruginosa*, regardless of the origin of the strains.

### Isolated SCV-like morphotype colonies are separated in two distinct clusters

By taking a closer look at the emerged SCV-like morphotypes, we observed that their sizes (Table 1) and overall appearance (Fig. 1) differ. Some colonies were denser, with well-defined round edges and others were more translucent with undefined edges (Fig. 1). We then asked whether these different types of SCV-like morphotypes are indeed *bona fide* SCVs, and if a distinction can be made between them. We focused on five strains representing the different origins, (Table 1, strains indicated by an asterisk) and isolated the various distinct morphotypic small colonies produced by each following static incubation and plating. Besides their sizes, we looked at several phenotypes typically associated with SCVs: swimming motility, biofilm formation and production of EPS, cell aggregation and production of c-di-GMP. Because cell aggregation induces the production of pyoverdine, the fluorescent siderophore of *P. aeruginosa*, while loss of the EPS coding genes, *pel* and *psl*, leads to inhibition of pyoverdine production (34), we used the production of pyoverdine as an indirect measurement of cell aggregation and EPS production. We compiled the phenotypical data for each distinct SCV-like morphotypes (SMs) (Table S1) and performed a principal coordinates analysis (PCoA) based on their colony size, auto-aggregation properties (pyoverdine production), their ability to perform swimming motility, timing of biofilm formation and density of biofilms. We found that the various distinct SMs generated by the five parental strains clustered in two separate groups (named Cluster 1 and Cluster 2) (Fig. 2). Members of both clusters for the SMs of soil strain 57RP, the sink hospital strain CL-511, the food strain PB PFR11 C2, and the clinical strain FC-AMT0134-9 had phenotypic features that distinguished them from their parental strain (Fig. 2). Cluster 2 of strain PA14 contained only one isolated SM, but we believe that this is only the result of lower abundance of this form when sampling was performed. These results indicate that two distinct phenotypic types of SCV-like morphotypes emerged in our culture conditions.

**Fig 2.**
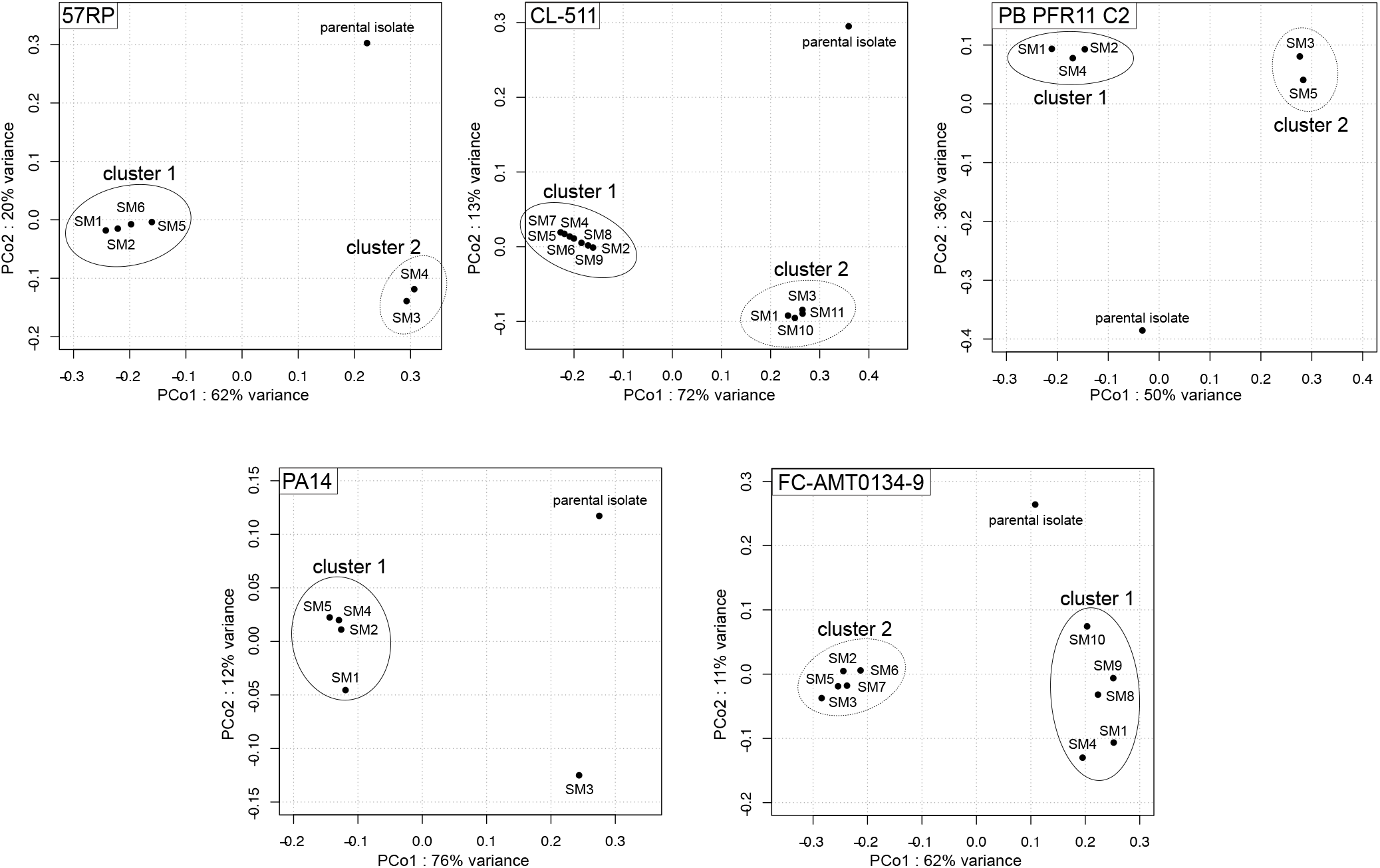
Small colonies isolated from static cultures are clustered in two separate groups according to their phenotypic features. PCoA analysis were performed with a matrix composed of data obtained from the phenotypic tests (swimming, biofilm formation, and pyoverdine production) for the parental strain and distinct small colonies isolated from static cultures with a diameter at least two times smaller than parental strain (Table S1). Each point represents a small colony isolated from the static cultures and have a name code composed of SMx standing for **<u>S</u>**mall **<u>M</u>**orphotype where x is anarbitrary number attributed during the isolation of the colonies. The identification of statistically distinctive clusters was performed using simprof tests and hclust.

### SMs from Cluster 1 are typical SCVs with a reversible state

SMs belonging to Cluster 1 of each strain share some common features: a reduced swimming motility, and/or a promoted biofilm formation, and/or enhanced auto-aggregation properties (pyoverdine production) compare with their parental strain (Table S1 and Fig. S1). These features are typical of SCVs described in the literature. Since these phenotypes are regulated by c-di-GMP, we assessed intracellular c-di-GMP levels in selected SMs of Cluster 1. As expected, higher c-di-GMP levels were measured in Cluster 1 SMs than in their parental counterparts, again indicating that Cluster 1 SMs are typical SCVs (Fig. 3). In addition to quantitative PCoA data, we looked at rugosity of SM colonies, a qualitative phenotype traditionally associated with SCVs. While Cluster 1 SMs colonies display a very distinctive rugose surface compared with their parental counterparts, rugosity appearance was diverse among the strains (Fig. 4).

**Fig 3.**
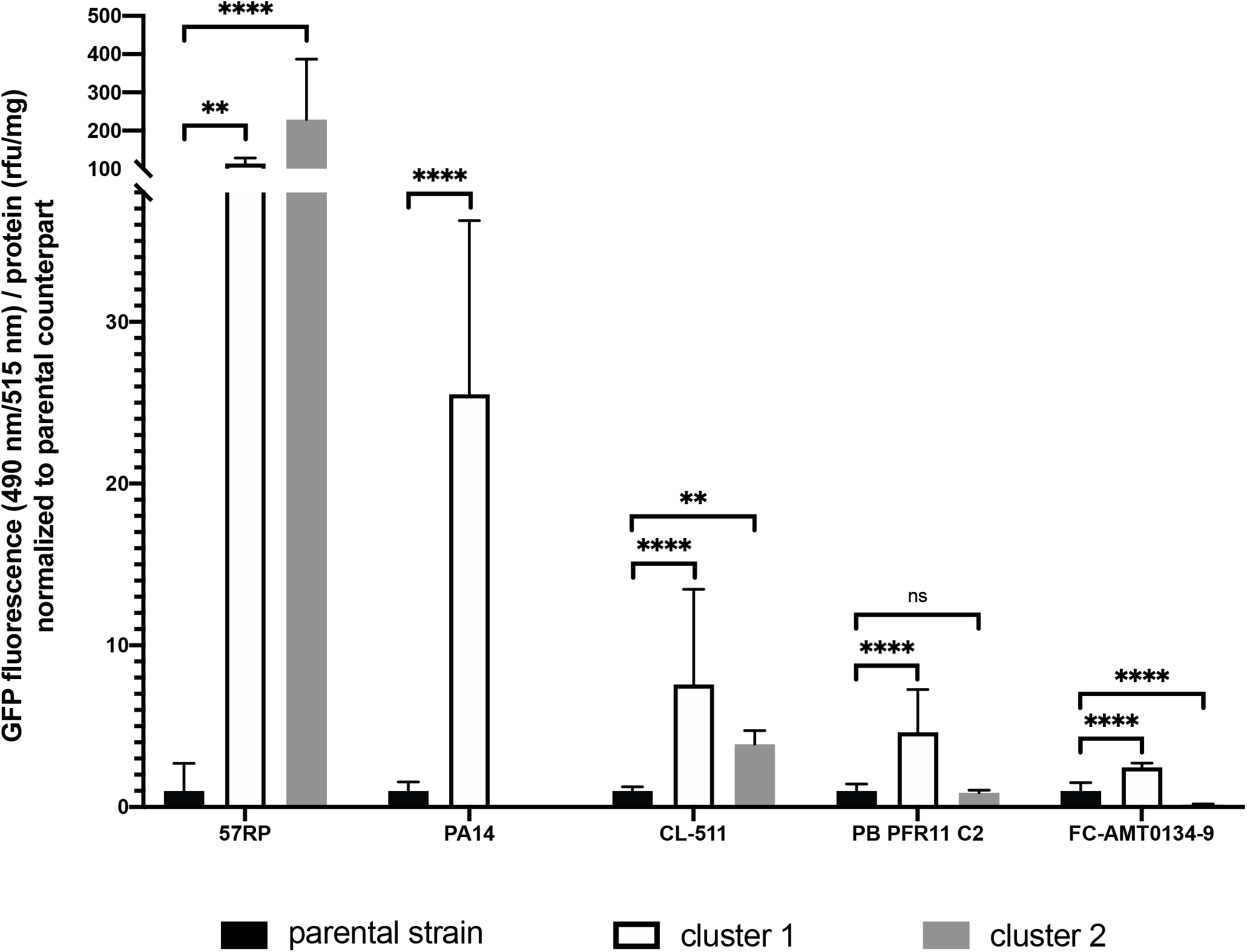
c-di-GMP production is altered for SMs from Cluster 1 and 2 compared with their respective parental strain. c-di-GMP production was measured with the fluorescent-based biosensor pCdrA-gfp on overnight washed cultures. The values are means standard deviations (error bars) for three transformants. Transformed morphotypes were SM2 and SM6 (cluster 1) and SM4 (cluster 2) for strain 57RP; SM4 and SM5 (cluster 1) for strain PA14; SM8 and SM9 (cluster 1) and SM10 (cluster 2) for strain CL-511; SM1 and SM2 (cluster 1) and SM3 and SM6 (cluster 2) for strain PB PFRC11 2; SM9 (cluster 1) and SM5 and SM7 (cluster 2) for strain FC-AMT0134-9. Stars represents the statistical significance of the results calculated by an Ordinary one-way analysis of variance (ANOVA), ****, P Value ≤ 0.0001; ***, P Value ≤ 0.001; **, P Value ≤ 0.01; *, P Value ≤ 0.05; ns, not significant. Data are normalized between them based on their parental strain.

**Fig 4.**
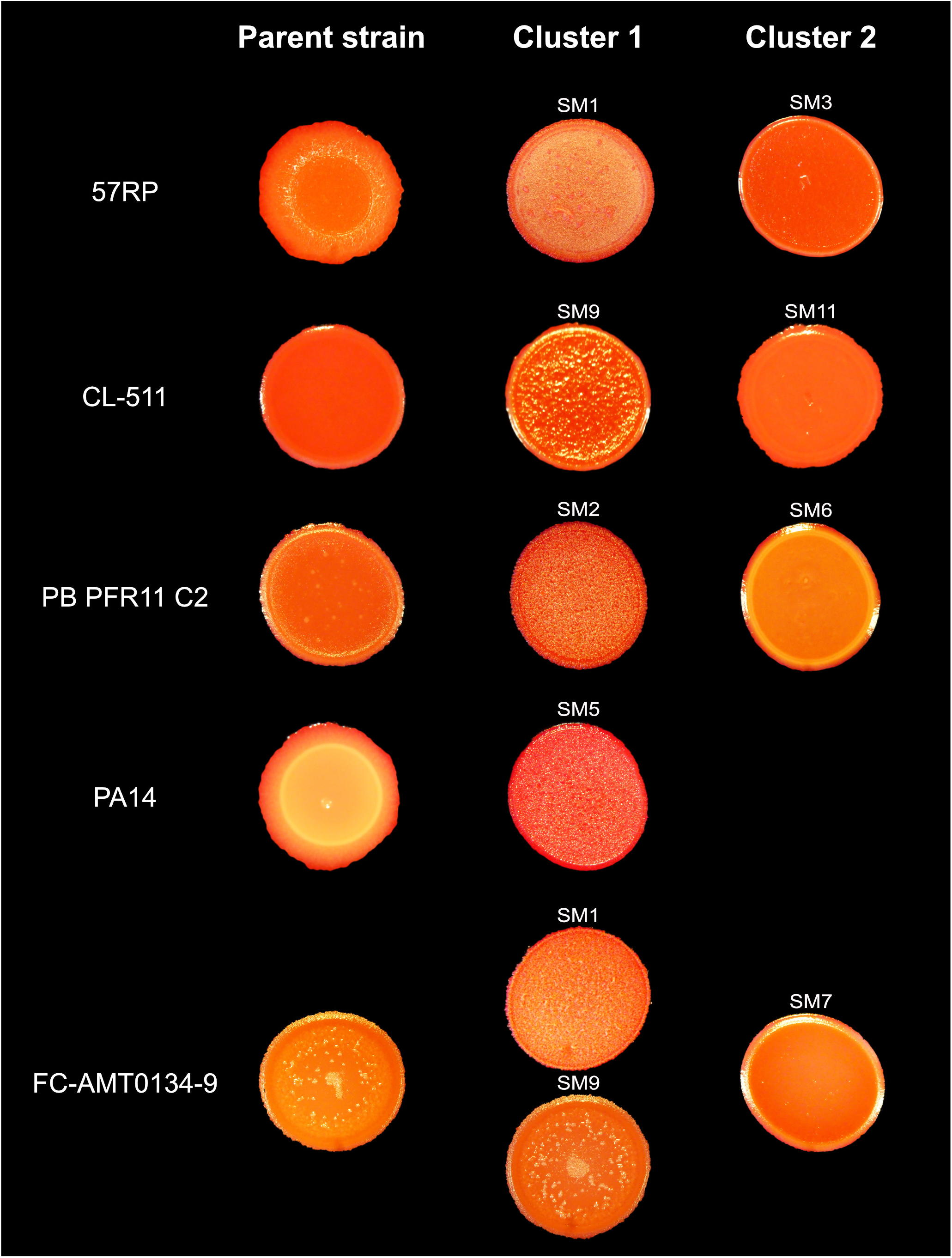
Appearance of colonies for the parental isolates and SMs from Cluster 1 and Cluster 2 on Congo Red plates. The SM showed for each cluster is representative of all the SMs included in one cluster since they have a similar appearance. Plates were observed with a binocular StemiDV4 (Zeiss) and photos were taken with a DMC-ZS60 camera (Panasonic Lumix), after 24 h of incubation at 30°C.

Finally, to further confirm that Cluster 1 SMs are indeed SCVs, we observed the expression of spontaneous reversion to a larger, parental-like phenotype, a property traditionally associated with phase variation. As stated above, SMs were readily obtained after a unique 65 h incubation under static culture conditions, suggesting that their emergence rate is high (Fig. 1). In addition, on agar plates, reversion to a parental-like morphotype was observed after a 48 h incubation at 30°C for SMs belonging to Cluster 1 (Fig. 5). Reversion was revealed as an outgrow from the original colony but sometimes observation was less evident, for instance in isolate PB PFR11 C2 reversion was revealed by an appearance change at the colony surface (Fig. 5). This reversibility, in addition to their phenotypical characterisation confirms that SMs from Cluster 1 are SCVs.

**Fig 5.**
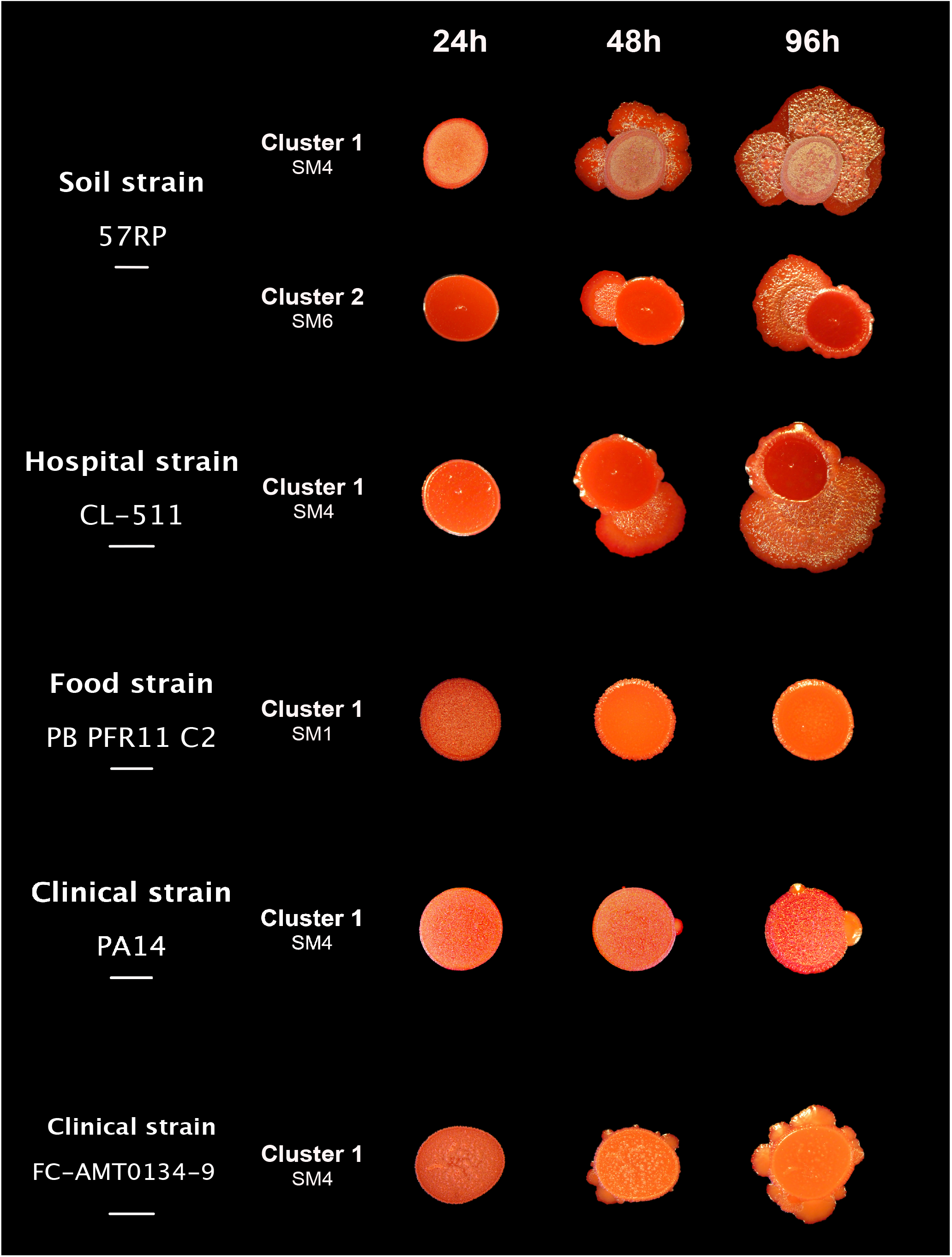
Reversion occurs on solid media for specific morphotypes after 48 h incubation. Ten µl of a culture of parental strain or a cluster representative morphotype (SMs) was dropped on 0.1% congo red TS-Agar 2% plates. Plates were observed with a binocular StemiDV4 (Zeiss) and photos were taken with the camera DMC-ZS60 (Panasonic Lumix), after 24 h, 48 h and 96 h of incubation at 30°C. Scale bars represent 5 mm.

### SMs from Cluster 2 display phenotypical heterogeneity

Unlike Cluster 1 SMs, SMs included in Cluster 2 display inter-strain diversity considering the phenotypes used for the PCoA (Table S1 and Fig. S1). For instance, Cluster 2 SMs swimming motility was intermediate between the parental strain and Cluster 1 SMs for strains 57RP and PB PFR11 C2 (Table S1 and Fig. S1, A). However, for strains CL-511 and FC-AMT0134-9 the swimming motility was increased compared to both Cluster 1 SMs and the parental strains (Table S1 and Fig. S1, A). In addition to PCoA data, c-di-GMP production in Cluster 2 SMs was also variable depending on the parental strain: 57RP Cluster 2 SMs showed higher levels of c-di-GMP compared with both parental strain and Cluster 1 SMs but CL-511 Cluster 2 SMs showed higher production of c-di-GMP only compared to the parental strain (Fig. 3). Also, Cluster 2 SMs in the food strain PB PFR11 C2 showed similar production of c-di-GMP and Cluster 2 SMs in the clinical strain FC-AMT0134-9 even lower production of c-di-GMP compare to their parental strain (Fig. 3). Thus, c-di-GMP levels are not a driving feature for SMs of Cluster 2. Colony surface aspects of Cluster 2 SMs on Congo Red plates was also distinct, once again depending on the parental strain. Colonies of SM3 and SM4 from 57RP displayed a rugose surface, however less pronounced than for Cluster 1 morphotypes (SM1, SM2, SM5 and SM6), in accordance with the reduced autoaggregative properties (Fig. 4 and Fig. S1, D). For the other strains (PA14, PB PFR11 C2, CL-511 and FC-AMT0134-9), SMs from Cluster 2 displayed a smoother surface on Congo Red, closer to the parental strain (Fig. 4). While we consider that Cluster 2 SMs are phase variants because of their rapid emergence to reproducible phenotypes, reversion to a larger colonial morphotype akin to WT was only observed for 57RP Cluster 2 SMs and not for the other strains, after 96 h (Fig. 5). All together, these results indicate that, apart from strain 57RP, SMs from Cluster 2 do not exhibit the majority of the traditionally described SCVs features.

## DISCUSSION

### Ability to switch to the SCV phenotype is a conserved feature among *P. aeruginosa* strains, regardless of their origin

SCVs have been reported several times in the context of human infections, notably in CF individuals. A correlation between the emergence of *P. aeruginosa* SCVs and infection persistence in animal models was established, supporting the idea that the SCV phenotype confers a fitness advantage under chronic infection conditions (35-37). Switch towards the SCV morphotype may represent an adaptation strategy to the hostile environment of the host by increasing resistance to host immunity and antimicrobial treatments (36, 38). However, the emergence of SCVs cannot be exclusively related to a clinical context. For instance, in 2001 Déziel *et. al*. (18) reported the emergence of SCVs in laboratory cultures of a soil *P. aeruginosa* isolate. However, since then, no SCVs have been reported from a non-clinical context, so the question of prevalence remained open: is the ability to adopt a SCV phenotype mostly restricted to clinical isolates, from chronic infections including a biofilm aspect, - or not?

Here, we investigated the distribution of a SCV-based adaptative strategy in *P. aeruginosa* by screening 22 strains from various origins. Screening was performed in static cultures, a growth condition that generates different microenvironments, as seen by the formation of a pellicle biofilm at the air-liquid interface. For all 22 strains, small colonies emerged in static cultures, with colonies isolated on agar plates with sizes similar to SCVs described in other studies (16, 18). However, SCVs are not exclusively defined by the smaller size of their colonies. SCVs are also often identified based on the rugosity of the colony formed on Congo Red agar plates. Indeed, SCVs are often referred as RSCVs for <u>R</u>ugose <u>S</u>mall <u>C</u>olony <u>V</u>ariant (14, 32, 36).Nevertheless, rugosity is a subjective feature, and its description may vary according to the observer and culture conditions. Indeed, we have observed that the rugosity level changes according to strains. This might be especially true for strains originating from various environments, as in the present study. Thus, we decided to take advantage of the various additional phenotypes described for SCVs to ascertain their identity. To this end, we focused on five strains representing diverse environmental origins. Based on their phenotypic features, the small colonies obtained from each parental strain were clustered into two distinct groups. Small colonies classified in Cluster 1 shared several inter-strain phenotypic features, including reversion after 48h. Based on what is already known on SCV characterisation, these small colonies can be defined as SCVs. This reveals that SCVs emerge from *P. aeruginosa* isolates from various origins. Thus, the ability to switch to the SCV phenotype is an intrinsic feature of the species.

### Switch to SCV is a reversible mechanism, likely to be regulated by phase variation through modulation of c-di-GMP

Phenotypic switching refers to a reversible interchange of states. Several studies suggest that phenotypic switching could be regulated by a reversible adaptation mechanism: phase variation (18, 39). Unlike reversible adaptation mechanism, genetic diversity generated by random mutations leads to a microbial subpopulation adapted to specific conditions. However, the acquired benefit will disappear when the environmental conditions fluctuate since genomes have been mutated irreversibly (19). Reversible adaptation mechanisms are based on DNA rearrangements and lead to variation in gene expression (19). Phase variation mechanisms lead to emergence of a heterogeneous population in which the best suitable phenotype will multiply until the conditions fluctuate again and the selected phenotypes revert to another phenotype. Phase variation is a common phenomenon in Gram-negative bacteria and is typical of bacteria thriving in heterogeneous ecological niches (20, 21, 40), notably *P. aeruginosa* (39). Indeed, phase variation mechanism represents a significant advantage for the rapid adaptation to sudden changes in the environment (41, 42). Interestingly, phenotypes traditionally related to SCV (motility, aggregation) are regulated by phase variation mechanisms (20). In addition, one recent study reports a large genomic inversion in *P. aeruginosa* SCVs (43). Thus, we hypothesize that the reversible switch to SCVs could be regulated by a phase variation mechanism. However, SCVs reversion can occur toward a phenotype likely different from the parental morphotype (22), suggesting that regulation is not necessarily an ON/OFF switch on a particular locus. It would be interested to investigate the ability of a revertant to switch again to the SCV phenotype under appropriate conditions. It should be emphasized here that colonies referred to as SCVs have been isolated from CF individuals and infected animals who actually had *wspF-*mutations (32), demonstrating that small colonies akin to SCVs can result from mutations and not phase variation.

Intracellular c-di-GMP levels regulate all of the phenotypes associated with SCVs: EPS production, motility, adherence, etc. (27-29). The c-di-GMP pool is regulated by diguanylate cyclases (DGC, synthesis of c-di-GMP) and phosphodiesterases (PDE, degradation of c-di-GMP) (44). In addition, emergence of SCVs can be “artificially” stimulated by introducing mutations in key genes involved in c-di-GMP regulation, such as the inhibitors coding genes *wspF* or *yfibNR* (14, 36, 45) or by overexpressing the DGC coding gene *wspR* (38). The phase variation mechanism at play to generate SCVs could function through regulation of c-di-GMP.

### Phase variation represents a conserved mechanism for rapid adaptation and persistence of a *P. aeruginosa* population

To readily observe the rapid adaptive benefit of phase variation, we need culture conditions where there is a strong selective pressure to form a biofilm. Déziel *et al*. (18) grew *P. aeruginosa* on an extremely hydrophobic source of carbon, hexadecane, so that the only way to thrive was to grow directly attached to the substrate, thus the need for rapid biofilm formation. However, this selection method is restricted to strains expressing the potential for aliphatic alkane catabolism (46). Here, we needed a selective condition more widely amenable to a general screen. When growing in a standing culture, oxygen is rapidly depleted and forming a biofilm at the air-liquid interface becomes the best solution, readily available to any strain able to produce a biofilm. Accordingly, we found that SCVs emerged spontaneously in a static (standing) liquid culture. Supporting this model, supplementing cultures with an alternative electron acceptor, such nitrate (as KNO_3_), reduced the emergence of SCVs in PA14 (Fig. S2).

SCVs have always been isolated in biofilm-promoting conditions or from environments where biofilms thrive (16, 31, 33). SCVs are especially prone at adherence and biofilm formation (18, 23, 31). The attached mode of growth (biofilm) is a widespread lifestyle in all types of environments (47-49). Biofilms are protective barriers for their bacterial components in the environment: they increase tolerance to antimicrobials such as antibiotics, disinfectants, toxic metals compared with free-living bacterial cells and they enhanced ability to survive in extreme conditions as instance desiccation (50-52). Thus, one can easily conceive that the switch to the SCV phenotype confers a significant advantage for colonization of various ecological niches, accounting for the conservation of the SCV phenotypic switch mechanism in all the tested strains. However, the exact link between SCVs and biofilm formation remains unclear; it is likely mostly relevant for the initial attachment to the surface/interface.

### Small colonies are not necessarily SCVs, nor variants

During our experiments with static cultures, we observed several small colony morphotypes Based on our PCoA analysis a proportion of them were clustered in two distinct groups (Fig. 2). Except for strain 57RP, the SMs from Cluster 2 did not display clear reversion after 48 h on solid medium (data not shown). However, SMs from Cluster 2 could still be able to revert in conditions outside the ones tested in our study. Also, their frequency of emergence seemed too high for mutants. Thus, we wonder if cluster 2 SMs should be identified as variants based on our criteria.

In contrast with SMs from Cluster 1, SMs from Cluster 2 showed inter-strain heterogeneous features. One hypothesis is that they represent intermediate forms between a SCV-like phenotype and reversion. Supporting this hypothesis, we observed a large diversity of morphotypes on plates prepared from our static cultures. Among them, large colonies also displayed features similar to revertants (16). This observation supports our hypothesis that reversion could have occurred in the static liquid cultures, and intermediate forms could consequently be isolated. Maybe several mechanisms can act in parallel to induce the phenotypic diversity we observed, thus increasing the likelihood that the best adapted subpopulation would be readily available to allow survival of the group.

The SCV phenotype has been linked to the persistence of *P. aeruginosa* in the context of infections in a human host, notably linked to its increased resistance against antimicrobials and host immunity. However, we have demonstrated that strains isolated from soil, food and hospital environments can also adopt a SCVs phenotype. This indicates that the ability of *P. aeruginosa* to form SCVs is naturally widespread, and SCVs emergence is not exclusively related to the pressure of the clinical environment. This is the first report of high prevalence of SCVs among *P. aeruginosa* strains, regardless of the origin of the isolates. The SCVs that were identified showed reversion after 48 hours on solid media. This result supports the hypothesis that *P. aeruginosa* uses a reversible adaptation strategy, generating phenotypic diversity, to rapidly adapt and persist into diverse environmental conditions, accounting for its versatility and persistence in a lot of environments. A deeper comprehension of the adaptation strategy used by *P. aeruginosa* could ultimately provide innovative strategies for eradication of this opportunistic multiresistant pathogen of public concern.

## MATERIALS AND METHODS

### Bacterial strains ang growth conditions

Bacterial strains are listed in Table 1 and their specific origin are listed in Table S2. In this study, the term “parental strain” designs the original strain used to evolve other morphotypes in static cultures, including SCVs. Strains were grown in tryptic soy broth (TSB; BD), at 37°C in a TC-7 roller drum (NB) at 240 rpm for the parental strains and at 30°C in an Infors incubator (Multitron Pro) at 180 rpm (angled tubes) for the isolated evolved morphotypes. Static cultures were inoculated with the parental strain at an initial OD_600_ of 0.05 and incubated at 30, 30.9, 32.2, 33.9, 36.3, 38, or 40°C for 65 hours. Cultures were then spread on tryptic soy agar 2% plates (TS-Agar; AlphaBiosciences) unless stated otherwise. Two percent agar were added to limit expansion of colonies and improve isolation of the distinct morphotypes.

### Bradford protein assay

Due to the highly aggregative properties of SCVs, OD_600_ measurements were not appropriate to evaluate growth of some of the isolated evolved morphotypes. The Bradford protein assay was used to quantify the concentration of total proteins in all our samples. Pellets from 1 ml of culture were resuspended in 1 ml 0.1 N NaOH and incubated 1 h at 70°C. Protein concentrations were measured on samples according to the manufacturer guidelines for the Bradford reagent (Alfa Aesar).

### Phenotypic tests

Overnight (O/N) cultures of parental strains and their isolated morphotypes were grown at 30°C in an Infors incubator (Multitron Pro) at 180 rpm in angled tubes. Since biofilms formation occurred in cultures, they were transferred to clean tubes before using to perform experiments or Bradford protein quantifications. Statistical analyses were achieved using Ordinary one-way analysis of variance (ANOVA). Each phenotypic test was performed in technical triplicates.

### Morphology on Congo red plates

A 1% Congo red solution in water (Fisher scientific) was added to TS-Agar 2% to a final concentration of 0.1%. Ten µL of culture were spotted on the plates. Plates were incubated at 30°C and observed after 24 h, 48 h and 96 h. Plates were observed with a binocular StemiDV4 (Zeiss) and photos were taken with the camera DMC-ZS60 (Panasomic Lumix).

### Swimming motility tests

Swim plates were prepared and dried for 15 min under the flow of a Biosafety Cabinet (20 mM NH_4_Cl, 12 mM Na_2_HPO_4_, 22 mM KH_2_PO_4_, 8.6 mM NaCl, 0.5% Casamino acids (CAA), 0.3% Bacto-Agar (BD), supplemented with 1 mM MgSO_4_, 1 mM CaCl_2_ and 11 mM dextrose). A volume of 2.5 µL of culture was inoculated in the agar. Plates were incubated 20 hours at 30°C. Swimming ability was assessed by measuring the area (mm^2^) of the turbid circular zone using ImageJ. All experiments were performed in triplicates.

### Biofilm formation

Microtiter (96-well) plates containing 1/10 TSB supplemented with 0.5% CAA were inoculated from a transferred overnight culture in order to obtain a starting concentration of 70 mM proteins. Each sample was inoculated in five different wells. Plates were incubated at 30°C without agitation. After 6 and 24 h, plates were rinsed thoroughly with distilled water and 200 µL of a 1% Crystal violet solution was added to each well. After 15 minutes of incubation at room temperature, plates were rinsed thoroughly with distilled water and the dye was solubilized in 300 µL in 30% acetic acid. The absorbance was measured at 595 nm with a microplate reader (Cytation3, Biotek). Bovine serum albumin (BSA) was used to generate a standard curve. Earliness of biofilm formation was calculated as the % of biofilm formed after 6 h of incubation compared with total biofilm formed after 24 h incubation. Density of the biofilm was calculated as the amount of biofilm formed after 24h.

### Pyoverdine production

Overproduction of pyoverdine was previously noted as a feature of strain 57RP SCVs (18). We confirmed that a SCV from PA14 showed high fluorescence level at pyoverdine wavelength, likely to account for cell aggregation and EPS overproduction. An SCV isolated from a PA14 *pvdD* mutant, which is no longer able to produce pyoverdine, showed lower fluorescence levels, similar to parental colonies, confirming that (1) pyoverdine production is responsible for the fluorescence detected and (2) measured fluorescence is correlated with SCV aggregation properties (Fig. S3). To measure pyoverdine production, black 96-well plates (Greiner) were filled with 200 µL of culture. Fluorescence was measured at wavelengths 390nm/530nm excitation/emission using a microplate reader (Cytation3, Biotek).

### C-di-GMP quantification

C-di-GMP levels were assessed with the fluorescence-based biosensor pCdrA-gfpC (53, 54). pCdrA-gfpC was constructed by Tim Tolker-Nielsen (addgene plasmid #111614; http://n2t.net/addgene:111614 ; RRID:Addgene_111614). Purified plasmids were transformed by electroporation in evolved morphotypes obtained from static cultures (55). Transformants were selected on TS-Agar 2% supplemented with 100 µg/ml gentamycin. Three clones for each transformed morphotypes were cultured in TSB supplemented with gentamycin 100 µg/ml. Cultures were washed twice in fresh TSB to get rid of a potential non-specific fluorescence due to secreted fluorescent pigments as pyoverdine. Fluorescence was measured using a Cytation3 microplate reader (BioTek) at 490nm/515nm (excitation/emission) in black 96-well plates (Greiner). The non-transformed strain was used as a control. Fluorescence from the control was subtracted to the fluorescence signal for the transformed strains.

### PCoA analysis

Colonies identified as SMs compared with their parental isolate (cf. results) were used to perform a principal coordinate analysis (PCoA). Statistical analyses were performed using RStudio software version 1.3.1093 (56) with normalised data showed in Table S1. A Euclidean distance matrix was used to generate a clustering of the bacterial isolates according to their phenotypical profile. A Similarity Profile Analysis (simprof) was performed to determine the number of significant clusters produced using hclust with the assumption of no a priori groups. Significant clusters were considerate when at least two evolved morphotypes constituted it.

## ACKNOWLEDGMENTS

We thank Cynthia Bérubé for her help with the c-di-GMP biosensor preliminary experiments, and Thays de Oliveira Pereira for critical reading of the manuscript.

This work was supported by grant MOP-142466 from the Canadian Institutes of Health Research (CIHR). Dr. Alison Besse is a Fellow of the postdoctoral grant Calmette and Yersin from the Institut Pasteur.

The funders had no role in study design, data collection and interpretation, or the decision to submit the work for publication.

AB, MCG, ED conceived the project, contributed to experimental design and interpreted results. AB and MT contributed to data acquisition. AB, MCG and ED wrote, reviewed and edited the manuscript.

